# Muscle synergies are flexibly recruited during gait pattern exploration using motor control-based biofeedback

**DOI:** 10.1101/2022.07.25.501482

**Authors:** Alyssa M. Spomer, Robin Z. Yan, Michael H. Schwartz, Katherine M. Steele

## Abstract

Understanding how the central nervous system coordinates diverse motor outputs has been a topic of extensive investigation. While it is generally accepted that a small set of synergies underlies many common activities, such as walking, whether synergies are equally robust across a broader array of gait patterns or can be flexibly modified remains unclear. Here, we evaluated the extent to which synergies changed as nondisabled adults (n = 14) explored gait patterns using custom biofeedback. Secondarily, we used Bayesian Additive Regression Trees to identify factors which were predictive of synergy modulation. Participants performed 41.1 ± 8.0 gait patterns using biofeedback, during which synergy recruitment changed depending on the type and magnitude of gait pattern modification. Specifically, a consistent set of synergies was recruited to accommodate small deviations from baseline, but additional synergies emerged for larger gait changes. Synergy complexity was similarly modulated; complexity decreased for 82.6% of the attempted gait patterns, however, distal gait mechanics were highly predictive of these changes. In particular, greater ankle dorsiflexion moments and knee flexion through stance, as well as greater knee extension moments at initial contact corresponded to a reduction in synergy complexity. Taken together, these results suggest that the central nervous system preferentially adopts a low-dimensional, largely invariant control strategy, but can modify that strategy to produce diverse gait patterns. Beyond improving understanding of how synergies are recruited during gait, study outcomes may also help identify parameters that can be targeted with interventions to alter synergies and improve motor control following neurological injury.

## 1. INTRODUCTION

Humans are capable of producing a broad array of movements, allowing for robust locomotion in diverse and unpredictable environments. To achieve this range of motor outputs, it has been hypothesized that the central nervous system (CNS) recruits a small number of synergies (i.e., modes, modules), defined as groups of coactivating muscles; this architecture is believed to simplify control beyond activating muscles independently (Bizzi and Cheung, 2013; Ting et al., 2015; Tresch and Jarc, 2009). Numerous studies have evaluated this hypothesis experimentally, employing matrix decomposition techniques, such as non-negative matrix factorization, to extract synergies and their corresponding activation patterns from electromyography (EMG) data (Lee, 1999; Tresch, 2005). These studies revealed that tasks such as walking (Allen and Neptune, 2012; Ivanenko et al., 2004), running (Cappellini et al., 2006; Hagio et al., 2015) and cycling (Barroso et al., 2014) share a small set of muscle synergies, despite being biomechanically distinct. Further, across tasks, changes in speed (Rozumalski et al., 2017), incline (Ivanenko et al., 2004; Rozumalski et al., 2017), cadence (Rouston et al., 2014), and body-weight loading (Ivanenko et al., 2004; McGowan et al., 2010) are shown to shift the phase or duration of synergy activations rather than the structure of the synergies themselves. These observations suggest that modest changes in sensory input or biomechanical demand are accommodated by altering the activation of invariant synergies and lend credence to their centralized role in coordination (Cheung et al., 2005; Torres-Oviedo and Ting, 2010). However, whether synergies are equally robust across a greater subset of achievable gait patterns or can be actively modified during gait is largely unknown (Jason J. Kutch and Valero-Cuevas, 2012; Tresch and Jarc, 2009).

Because synergies generally align with the sub-tasks of walking (e.g., push-off, weight acceptance), gait patterns which impose additional mechanical requirements or present large changes in somatosensory feedback may alter synergy recruitment (Cheung et al., 2005; Ivanenko et al., 2005; Nazifi et al., 2017; Torres-Oviedo and Ting, 2010). This is supported by prior work in animal models which demonstrated that frogs recruit task-specific synergies during swimming, jumping, and walking which correspond to the unique biomechanical demands of each movement (d’Avella and Bizzi, 2005). Similarly, humans recruit specific synergies during perturbation recovery tasks to maintain mediolateral stability and reduce center of mass movement (Krishnamoorthy et al., 2004; Martino et al., 2015; Nazifi et al., 2017; Torres-Oviedo and Ting, 2010). Importantly, such synergies emerge in addition to those shared with other tasks, which suggests that the CNS flexibly draws from a limited library rather than deploying unique control strategies to accommodate task demand (Torres-Oviedo and Ting, 2010).

Taken together, prior results indicate that a relationship exists between the biomechanical constraints of a given task and the recruited control strategy. That is, the CNS may preferentially tune the activation timing of a consistent set of synergies but is simultaneously capable of recruiting different synergies to produce diverse outputs. Understanding when and how synergies are modulated across changing biomechanical contexts and the factors driving this modulation is critical to better inform how the CNS coordinates complex movement. While this relationship has been previously characterized across broad balance (Torres-Oviedo and Ting, 2010) and finger force generation tasks (Jason J Kutch and Valero-Cuevas, 2012; Valero-Cuevas et al., 2009), gait has not been studied to the same extent (Rouston et al., 2014; Zelik et al., 2014).

Beyond enhancing understanding of the neural control of gait, characterizing whether synergies can be modulated in walking may also inform methods for targeting aberrant synergy recruitment. Individuals with cerebral palsy (Schwartz et al., 2016; Steele et al., 2015; Tang et al., 2015), Parkinson’ s disease (Rodriguez et al., 2013), and spinal cord injury (Fox et al., 2013) as well as stroke survivors (Cheung et al., 2012; Clark et al., 2010) recruit fewer synergies than nondisabled peers which impacts independent mobility (Bowden et al., 2010; Clark et al., 2010; Mehrabi et al., 2019) and may reduce the efficacy of traditional interventions (Schwartz et al., 2016). Because available interventions for these populations often fail to alter synergies (Shuman et al., 2019), developing new paradigms to directly improve synergy recruitment has become a critical priority in gait rehabilitation. This has spurred the development of biofeedback and robotic gait training paradigms which have thus far yielded promising, yet still highly variable results (Booth et al., 2019; Conner et al., 2021; Rouston et al., 2013). As such, mapping the relationship between biomechanical constraints and synergy modulation may further inform the design of these systems by highlighting gait parameters that can be directly targeted to produce greater and more consistent changes in motor control.

The aim of this study was to characterize the robustness of synergies to changing biomechanical constraints during walking. Specifically, we evaluated the extent to which nondisabled individuals could modulate both synergy structure and complexity during walking while using motor control biofeedback to drive broad gait pattern exploration. These data were then used to build a Bayesian Additive Regression Trees (BART) model to identify biomechanical variables that were predictive of synergy modulation. We hypothesized that changing biomechanical constraints would alter the recruitment but not the structure of muscle synergies, but that different synergies may be recruited to accommodate large deviations from baseline. The results from this investigation will provide further insight into the extent to which motor control can be altered and, importantly, improve understanding of how the CNS shapes its control strategy to produce a repertoire of motor outputs. The latter will support the development of intervention strategies to improve motor control among individuals with neurological injury.

## 2. METHODS

### 2.1 Experimental Protocol

Fourteen nondisabled individuals (7M/7F; Age: 24.1 ± 4.7 years; Height: 1.7 ± 0.1 m; Mass: 65.7 ± 20.1 kg) were recruited to evaluate synergies during gait pattern exploration. Prior to participation, all provided written informed consent and the experimental protocol was approved by the University of Washington Institutional Review Board.

Participants walked on a treadmill at a self-selected speed (1.07 ± 0.13 m/s; Bertec, Columbus, OH) while responding to a custom biofeedback system, designed to encourage gait pattern exploration. Briefly, this system presented the participant with a real-time score of their dominant-limb synergy complexity, defined as the total variance accounted for by one synergy, on a graphical display (Steele et al., 2015). To facilitate participant interpretation, the displayed score was normalized to baseline walking and scaled such that a value of 100 corresponded to baseline and higher values indicated more complex control (see S1 for additional system details). Participants performed one baseline walking trial with the feedback system turned off followed by feedback trials during which they were instructed to either (1) raise or (2) lower their complexity score; two trials were performed in each target direction. All trials were three minutes long and separated by mandatory one-minute rest periods. During the feedback trials, participants were encouraged to explore a broad range of gait patterns to modify their score. The only imposed restrictions were that they must (1) maintain forward-facing walking and (2) take at least five consecutive strides in the pattern selected.

Surface EMG data (Delsys Inc, Natick, MA) were recorded bilaterally for seven lower limb muscles: gluteus maximus (GM), lateral hamstrings (LH), medial hamstrings (MH), vastus medialis (VM), soleus (SO), tibialis anterior (TA), and medial gastrocnemius (MG). Raw EMG signals were low passed filtered (4^th^ order Butterworth; 20 Hz), rectified, and high pass filtered (4^th^ order Butterworth; 10 Hz) to establish linear envelopes (Shuman et al., 2017). After filtering, non-physiological signal spikes were removed using a robust-PCA algorithm (Lin et al., 2013) and the data were normalized to the 95^th^ percentile of maximum muscle activity across all trials.

Full-body motion data were collected using a 10-camera motion capture system (120 Hz) and a modified Helen Hayes marker set (Kadaba et al., 1990). Joint kinematics and kinetics were derived from marker data in OpenSim v3.3 using a 33 degree-of-freedom model, scaled to each subject (Delp et al., 2007; Rajagopal et al., 2016). The average root-mean-squared (RMS) and maximum error for all developed models were 1.3 cm and 2.5 cm, respectively, which fall below the recommended thresholds for model fidelity (Hicks et al., 2015).

### 2.2 Gait Analysis

Because participants explored many different gait patterns using the biofeedback system, we first had to extract each pattern attempted across trials and participants (Figure 1). To do this, the gait deviation index (GDI) was calculated from the kinematic data for every stride in each trial (Schwartz and Rozumalski, 2008). The GDI is a summary measure of deviations in pelvis, hip, knee, and ankle kinematics from ‘normative’ trends and was, therefore, expected to change during gait pattern exploration. For each trial, groups of five or more consecutive strides with similar GDI values were automatically labeled as unique gait patterns; each unique pattern identified was then subsequently confirmed via manual inspection to ensure appropriate labeling. Following labeling, average kinematic and kinetic trends at the pelvis, hip, knee, and ankle were quantified for each unique pattern. To identify kinematically-similar strategies adopted by multiple participants, the average kinematics for all unique patterns were separated into clusters (K_1_ to K_N_) using k-means clustering (Rozumalski and Schwartz, 2009).

**Figure 1:**
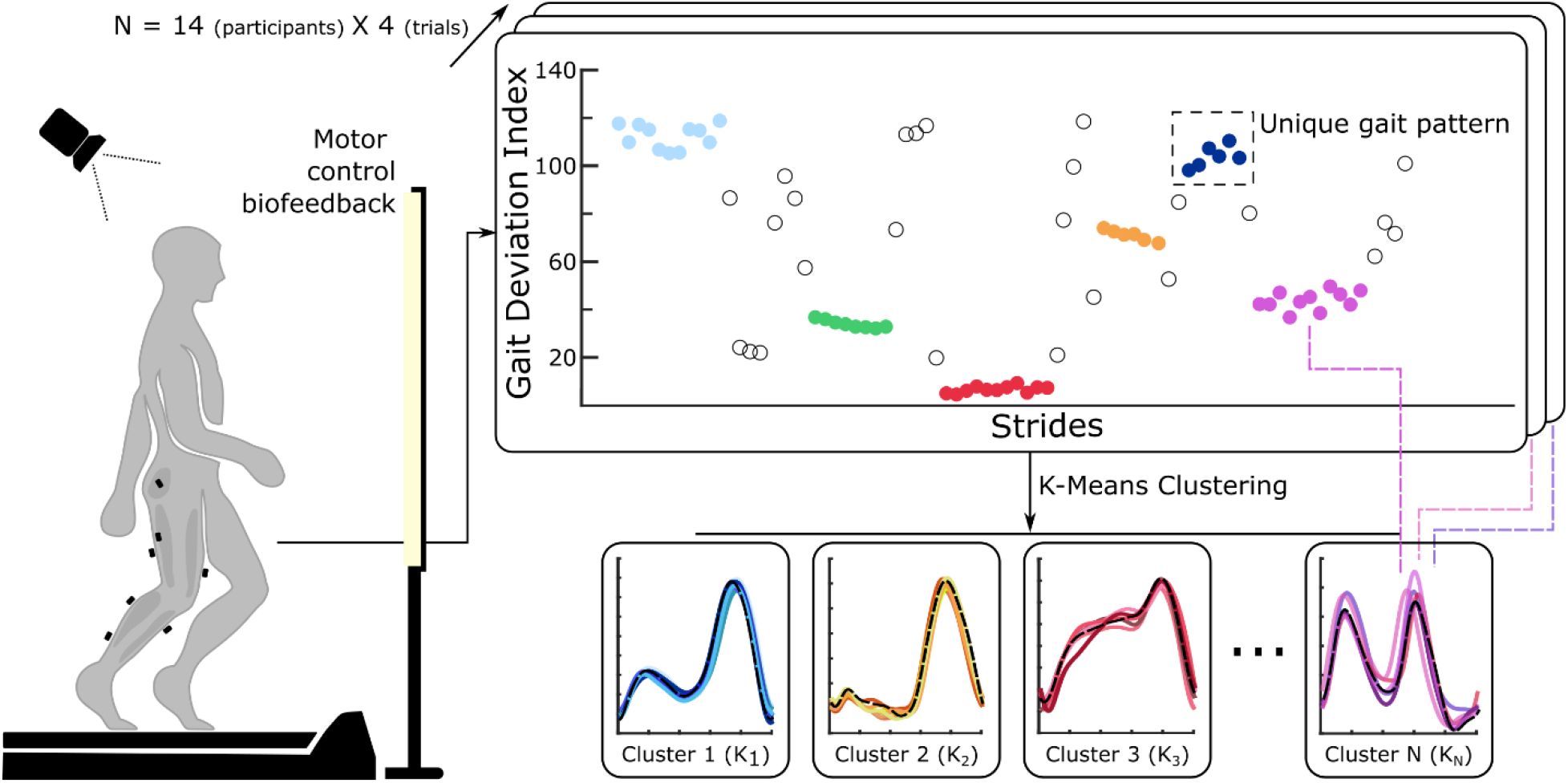
Methodology used to identify clusters (K_1_ to K_N_), representing kinematically-similar gait patterns attempted by participants during feedback walking. Full-body kinematic and kinetic data and lower-limb EMG data were collected while participants explored a broad range of movement patterns using biofeedback. The Gait Deviation Index (GDI) was calculated from kinematic data for each recorded stride for every participant and trial (56 data sets). Unique gait patterns were labeled as five or more consecutive strides with similar GDI values and manually confirmed. Kinematic data for these unique patterns were input into a k-means clustering algorithm to identify clusters (K1 to K_N_) across participants and trials.

### 2.3 Synergy Analysis

Muscle synergies were quantified from EMG data for each unique gait pattern using non-negative matrix factorization (NMF). NMF is a linear matrix decomposition technique which is commonly used to identify non-negative synergies (W) and their corresponding activations (C) from EMG data, such that *EMG*_*mxt*_ *= W*_*mxi*_*\*C*_*ixt*_ *+ error* where *m* is the number of muscles, *i* is the number of synergies, and *t* is the time points (Lee, 1999; Ting and Chvatal, 2010). The structure of the W and C matrices provide insight into how muscles coactivate across the gait cycle. Similarly, the total variance accounted for (tVAF) by a given number of synergies (*i*) can provide a summary measure of synergy complexity and has been frequently used as a marker for impairment level; individuals with neurological injury (Cheung et al., 2012; Clark et al., 2010; Fox et al., 2013; Rodriguez et al., 2013; Schwartz et al., 2016; Steele et al., 2015) have higher tVAF values (*i*.*e*. less complex control) for a given synergy solution (*i*) than nondisabled peers. Therefore, if synergies were sensitive to imposed biomechanical constraints, we may expect to see changes in both synergy structure and complexity measures.

We calculated *i* = 1 to 7 synergies using the dominant-limb EMG data for five concatenated strides for each unique gait pattern attempted. Because synergy analysis is sensitive to the amount of EMG data used, we elected to analyze a consistent number of strides across all patterns and participants (Oliveira et al., 2014). If a participant took more than five strides in a unique pattern, we performed a bootstrapping procedure by quantifying synergies using five random strides, selected with replacement from the available set, and replicating this process until a normal distribution was achieved; average synergies and tVAF values were then reported. The same bootstrapping procedure was performed on the baseline walking data with sets of five concatenated strides (replicates = 200) to ensure accurate comparisons between baseline and feedback conditions.

We evaluated synergy structure during gait pattern exploration in two ways. We first compared the inter-cluster (K1 to KN) similarity of synergy weights (W) and activation patterns (C) for the *i* = 3 synergy solution. This solution was evaluated, as three synergies explained over 90% of the variance in EMG data for the majority of unique gait patterns. We sorted synergy weights for all unique gait patterns attempted during exploration as well as baseline walking into *k* clusters (MacQueen, 1967). Because individuals may recruit different synergies during exploration compared to baseline gait, we varied *k* between *k* = *3* (i.e., synergies were consistent between baseline and exploration) and *k* = 10\**3* (*i*.*e*., different synergies emerged during exploration) and selected *k* as the number of clusters with the maximum silhouette coefficient (Rousseeuw, 1987); the upper bound on *k* was highly conservative and based on our expectation that synergies would be predominantly shared across gait patterns (Torres-Oviedo and Ting, 2010). Synergy weights and activations for each unique gait pattern were then sorted into their respective clusters (K1 to KN) and the average values were calculated. Secondarily, we evaluated the intrasubject similarity of baseline synergies with those recruited during exploration. This was done by fixing the W matrix as the synergy weights extracted from baseline walking for the three-synergy solution (*i = 3*) and using the multiplicative update rule from NMF to find a C matrix which minimized the error between W*C and the EMG data for each unique gait pattern that an individual attempted. From this, we were able to calculate the total variance that could be explained in each unique gait pattern by baseline weights (tVAF_3_BASE_) which was then compared to the tVAF_3_ values (i.e., those calculated directly from the EMG data for each unique gait pattern), yielding a measure of synergy similarity. If similar synergies were recruited during gait pattern exploration and baseline walking, we would expect tVAF_3_BASE_ and tVAF_3_ to be similar.

### 2.4 Statistical Analysis

#### 2.4.1 Cluster-wise comparisons

For each cluster (K_1_ to K_N_), we compared mean tVAF values to (1) baseline walking and (2) tVAF_BASE_ using paired t-tests to evaluate if synergy complexity or structure changed during gait pattern exploration, respectively. Secondarily, one-way ANOVA tests were used to compare if synergy complexity and structure were similar between clusters (K_1_ to K_N_); for any test that reached significance, t-tests were used to perform pairwise comparisons. Average kinematic trends at key phases within the gait cycle (*e*.*g*., push-off, initial contact) for each cluster (K_1_ to K_N_) were also compared to baseline walking using paired t-tests. To characterize stride-to-stride variability, the standard deviation of each kinematic parameter during exploration was also compared to baseline using paired t-tests. For all comparisons to baseline walking and post-hoc analyses, p-values were adjusted using a Holm-Šídák correction to account for multiple tests. We defined significance as p < α for α = 0.05 and report mean values ± 1 SD unless otherwise indicated. All cluster-wise statistical analysis was performed using the MATLAB Statistical Toolbox (MathWorks, Natick, USA).

#### 2.4.2. BART analysis

To further examine the relationship between gait pattern exploration and synergy complexity, we developed a Bayesian Additive Regression Trees (BART) statistical model (Chipman et al., 2010). BART is a ‘sum-of-trees’ machine learning method used for non-parametric function estimation, similar to other techniques such as boosting (Freund and Schapire, 1997; Schapire, 1990) and random forests (Breiman, 2001). However, unlike other methods, BART uses a regularization prior to control tree depth and shrinkage, effectively constraining individual trees as ‘weak learners’ to prevent data overfitting (Chipman et al., 2010; Kapelner and Bleich, 2016). BART was selected for this application due to its favorable predictive performance compared to other machine learning algorithms and because it can capture the non-linear relationships inherent in motion data (Chipman et al., 2010; Dorie et al., 2019; Tan and Roy, 2019).

We developed a BART model to predict changes in synergy complexity during exploration compared to baseline walking, quantified as the difference in total variance accounted for by a one-synergy solution (i.e., ΔtVAF_1_). Our predictor set (Table 1) included kinematic and kinetic variables that characterized each unique gait pattern as well as other metrics which could influence the type of gait patterns a participant attempted. When defining kinematic and kinetic predictor variables, we prioritized a set that captured salient trends at the pelvis, hip, knee, and ankle, while simultaneously maintaining predictor set conciseness. These criteria resulted in the variables outlined in Figure 2 (n = 31). For each of the identified kinematic and kinetic variables, both the mean and standard deviation values are included in the predictor set, normalized to baseline walking. We elected to include standard deviation measures in the model, as tVAF_1_ is sensitive to the amount of variance in the data and could, therefore, be affected by individuals simply moving with greater stride-to-stride variability, as might be expected during novel gait pattern exploration (Sawers et al., 2015). We tuned hyperparameters for the developed BART model using 10-fold cross-validation (parameters: k = 3, q = 0.9, nu = 3, num_trees = 200, seed = 30) and report both pseudo-R^2^ and the out-of-sample root-mean-squared error (RMSE) as metrics of model quality.

**Table 1:** BART Model Variables

| Response | Description |
| --- | --- |
| $\Delta tVAF_1$ | Difference in motor control complexity between each unique gait pattern and baseline walking. |
| Predictors | Description |
| Baseline $tVAF_1$ | Measure of motor control complexity during the baseline walking condition. Values range from 0 - 1, where a higher value indicates less complex control. |
| Participant Number | Values range from 1-14. This variable was used to evaluate if participant-level differences in biofeedback exploration emerged. |
| Speed | Nondimensional walking speed normalized to participant leg length. |
| Kinematics/Kinetics | Mean values of all variables outlined in Figure 2. Variables input as z-scores, normalized to baseline walking. These variables were selected as they sufficiently capture kinematic/kinetic trends during the gait cycle. |
| Kinematic/Kinetic Variability | Difference in the standard deviation of all variables outlined in Figure 2 between each unique gait pattern and baseline walking. These variables reflect the participant's ability to consistently produce the attempted gait patterns. |

**Figure 2:**
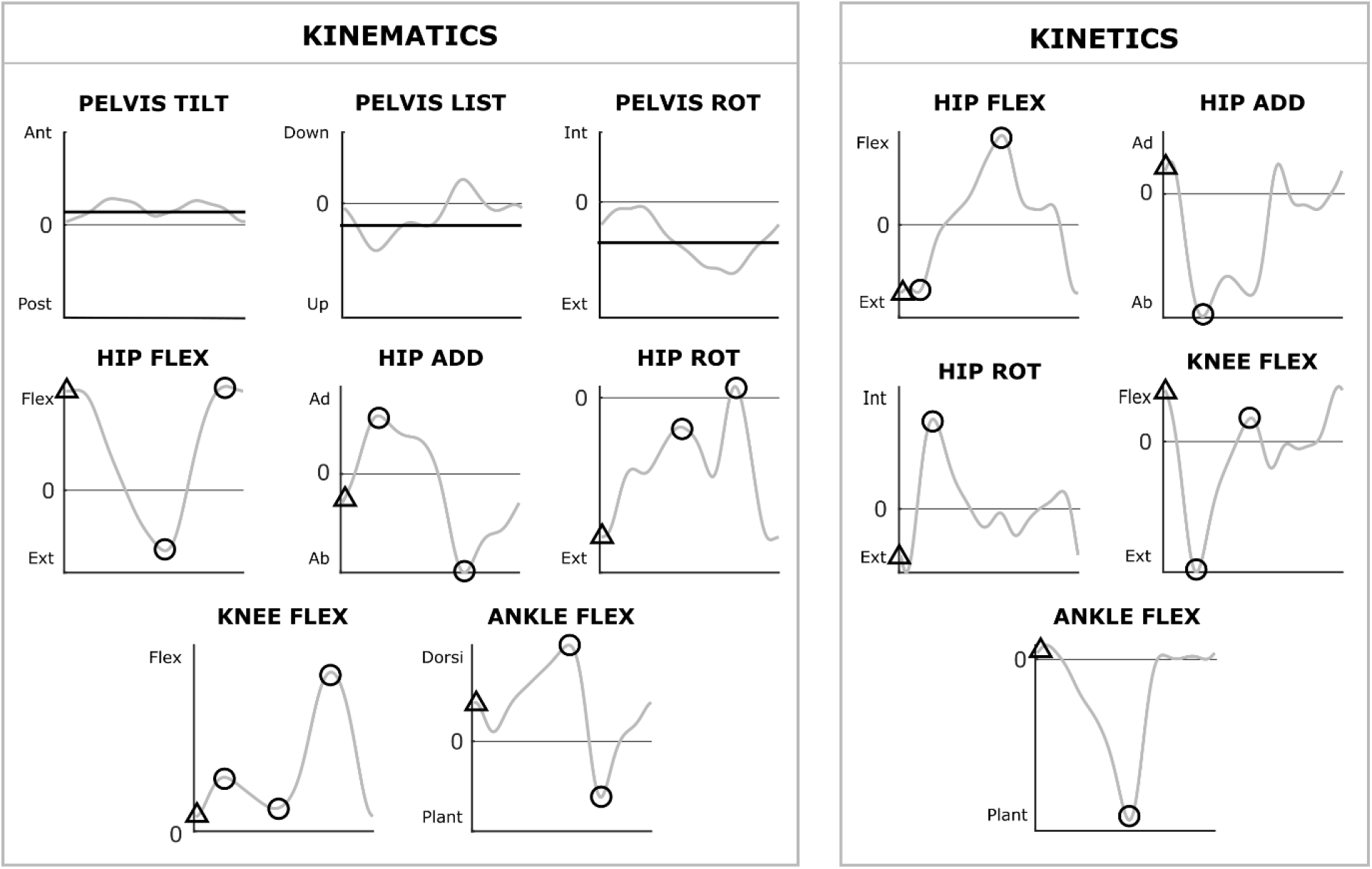
Kinematic and kinetic predictor variables included in the BART model. Each icon indicates a variable (n = 31 total) that was identified for every unique gait pattern. Variables were selected to capture trends at the pelvis, hip, knee, and ankle that could change during exploration. Circles indicate local maximum or minimum values, and triangles indicate initial contact points, calculated as the mean value over the first 5% of the gait cycle. Average pelvis list, tilt, and rotation angles across the gait cycle were included to capture asymmetries. The standard deviations of each kinematic and kinetic variable, used to capture stride-to-stride variability during gait pattern exploration, were also included in the predictor set.

Outputs from the BART model were interpreted with accumulated local effect (ALE) plots (Apley and Zhu, 2020). ALE plots are used to visualize the effect that individual predictors have on the specified response variable (i.e., ΔtVAF_1_), conditioned on all other model covariates. Unlike partial dependence plots, which are also commonly used, ALE plots are generated by averaging and accumulating the local rather than marginal effects of each predictor, making them unbiased in cases where predictors are highly correlated; this is particularly advantageous for this application, due to the high level of correlation between kinematic and kinetic variables during gait.

Because ALE plots are generated by sampling from the available data, some discrepancy between the ‘true’ and ‘estimated’ effect is expected (Apley and Zhu, 2020). To capture this uncertainty, we performed a boostrap analysis (n = 100 replicates), drawing samples with replacement from the original data set to generate a series of ALE plots from which the average and standard deviation could be quantified. Using these average plots, we approximated net effects for each predictor as the difference between the 95^th^ and 5^th^ percentile of the response. If synergies were sensitive to biomechanical constraints during gait pattern exploration, we would expect both kinematic and kinetic variables to have large net effects on ΔtVAF_1_. BART model development and ALE plot generation were performed in RStudio (RStudio Team, 2020) using the *bartMachineCV* and *ALEPlot* packages (Apley and Zhu, 2020; Kapelner and Bleich, 2016).

## 3. RESULTS

### 3.1 Gait Exploration

Participants explored 10.3 ± 2.8 unique gait patterns per feedback trial on average, resulting in 575 total patterns across all participants. These data were separated into five clusters, representing the common kinematic strategies attempted (Figure 3). K_2_ and K_4_ represented 24 and 78 unique gait patterns, respectively, and were characterized by increased hip flexion (K_2_: 47.7 ± 11.3°; K_4_: 23.1 ± 8.8°), knee flexion (K_2_: 70.6 ± 9.1°; K_4_: 40.9 ± 11.1°), hip abduction (K_2_: 7.4± 7.0°; K_4_: 7.7 ± 7.6°), anterior pelvic tilt (K_2_: 14.1 ± 6.6°; K_4_: 8.4 ± 7.9°), and ankle dorsiflexion (K_2_: 20.7 ± 2.7°; K_4_: 15.6 ± 5.2°) through stance compared to baseline. K_3_ represented 98 unique gait patterns defined by greater anterior pelvic tilt (2.8 ± 3.7°), hip abduction (3.9 ± 4.2°), and plantarflexion (6.3 ± 8.9°) through stance, as well as decreased knee flexion through swing (29.8 ± 9.1°). K_5_ included patterns with increased hip (44.7 ± 12.2°) and knee flexion (80.0 ± 10.1°) during swing and increased hip abduction (4.5 ± 3.5°) in stance. Finally, K_1_ included 322 unique gait patterns that aligned closely with baseline trends (p > 0.054 for all angles), capturing points within the feedback trials in which participants were minimally exploring.

**Figure 3:**
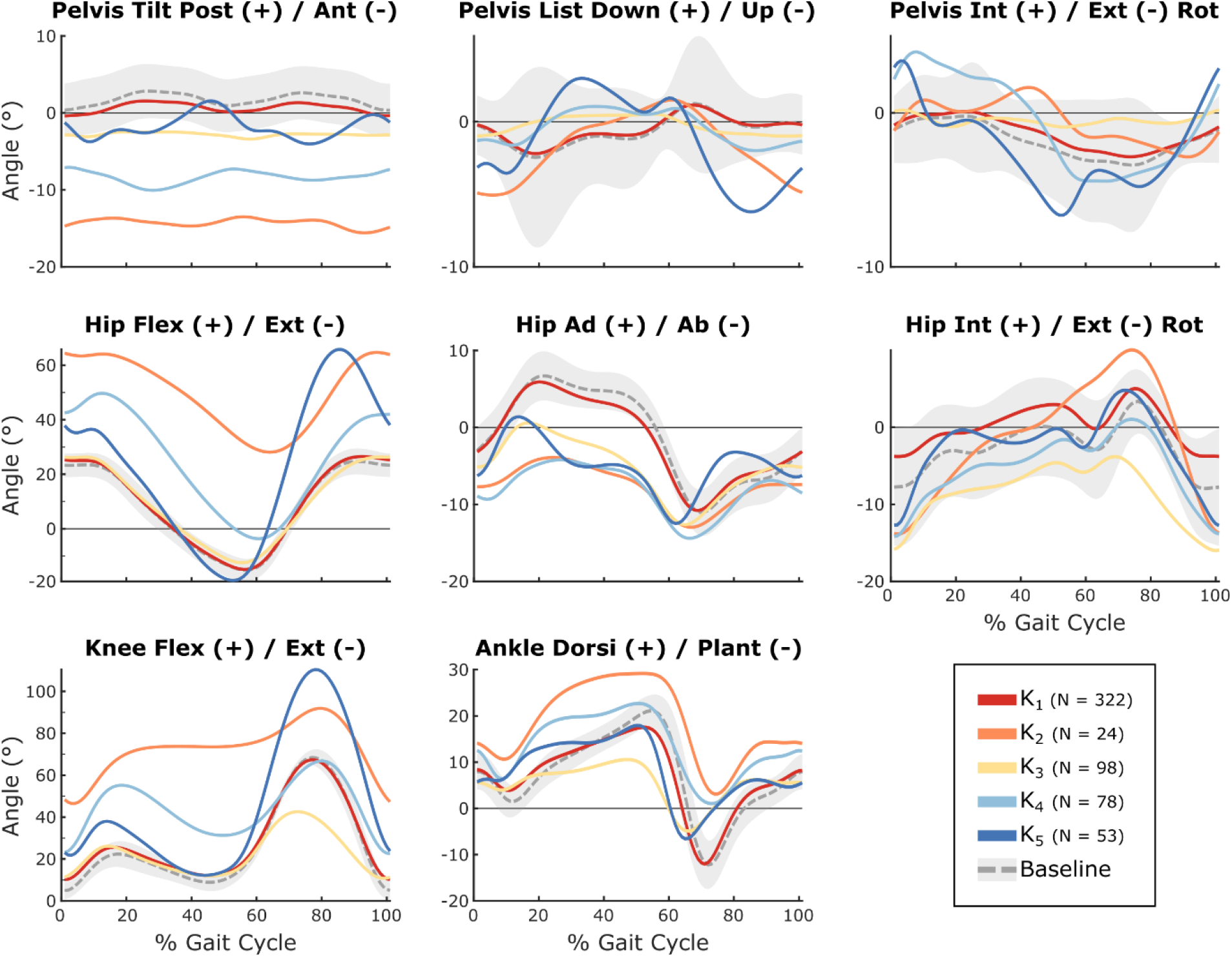
Average pelvis, hip, knee, and ankle kinematics for the five clusters identified by k-means clustering (K_1_ – K_5_), representing common gait patterns attempted during exploration. The baseline condition shows ± 1SD.

As expected, stride-to-stride variability increased for all kinematic parameters in K_2_ to K_5_ (p < 0.05 for all parameters), with the largest variability seen in gait patterns in K_2_. This increase in variability highlights an inherent learning effect associated with unique gait pattern reproduction. Even when participants were minimally exploring (*i*.*e*., K_1_), there was generally an increase in variability compared to baseline, likely due to the added attentional demand of responding to the biofeedback system.

### 3.2 Synergy Analysis

All participants were able to significantly modify synergy complexity during exploration (Figure 4). A one-synergy decomposition (*i* = 1) accounted for 66.1 ± 5.9% of the variance in the EMG data during baseline. When clustered, tVAF_1_ was 71.6 ± 7.2% (K_1_), 78.3 ± 6.6% (K_2_), 76.5 ± 6.4% (K_3_), 76.0 ± 6.2% (K_4_), and 69.8 ± 5.7% (K_5_), indicating that all of the explored patterns significantly decreased complexity (p < 0.05). Interestingly, there were also significant inter-cluster differences in tVAF_1_, suggesting that the type of gait pattern modification impacted complexity (p << 0.001). It should be noted that although participants were instructed to either raise or lower their synergy complexity scores, minimal differences existed between these trials; participants generally decreased complexity regardless of the target direction. As such, we did not conduct further analyses comparing target directions.

**Figure 4:**
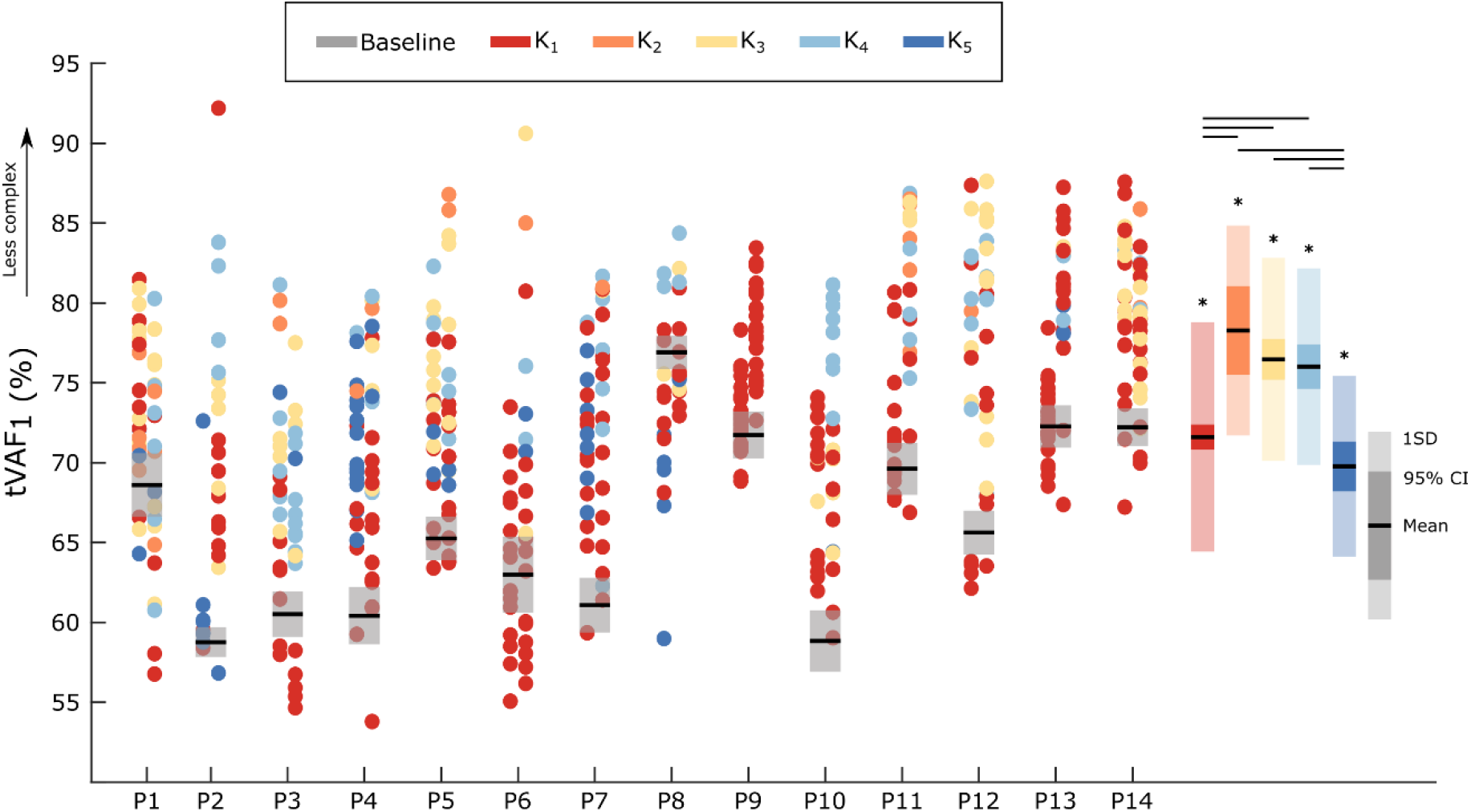
The total variance accounted for by a one-synergy solution (tVAF_1_) for each participant (P1-P14). Each dot represents a unique gait pattern and is colored according to the cluster it was sorted into (K_1_ to K_5_). For each participant, data is organized into two columns representing the feedback trials in which participants were instructed to decrease (left) and increase (right) their tVAF_1_. Baseline data is presented as a mean ± 1SD, representing the distribution of tVAF_1_ values resulting from bootstrapping using sets of five random strides (replicates = 200). Boxes represent the mean (black line), 95% confidence interval (solid color), and standard deviation (shading) of tVAF_1_ values for each cluster and the baseline condition. Larger tVAF_1_ values correspond to decreased motor control complexity. * denotes significant difference between each group and the baseline condition and black bars indicate significant inter-cluster differences (α = 0.05).

A three-synergy solution (*i* = 3) accounted for 92.6 ± 2.3% of the variance for all exploration and baseline walking patterns. Clustering yielded four distinct synergy structures (Figure 5) that were dominated by the TA (W_1_), hamstrings (W_2_), the quadriceps and gluteus maximus (W_3_), and the plantarflexors (W_4_). All four synergy structures were observed across K_1_ to K_5_ as well as baseline walking but were recruited with varying frequency. For example, baseline walking was primarily defined by W_2_, W_3_, and W_4_, which were present in 85.7%, 78.6%, and 100% of gait patterns in the group, respectively. These synergies align with those previously reported in nondisabled adults during steady-state walking (Allen and Neptune, 2012; Clark et al., 2010). In contrast, K3 was dominated by W1 (76.5%), W2 (82.7%), and W_4_ (91.8%). Interestingly, the plantarflexor synergy (W_4_) emerged for the majority of patterns in all clusters (K_1_ to K_5_) whereas W_1_, W_2_, and W_3_ were differentially recruited. These results suggest that a small pool of synergies exists that can be selectively drawn from depending on the biomechanical constraints of a given pattern. Across groups, synergy activation patterns were also distinct from baseline and aligned with key kinematic trends. For example, K_2_ was characterized by increased knee flexion and ankle dorsiflexion through the gait cycle, which was reflected in the increased activation of W_1_ in swing and W_3_ through stance.

**Figure 5:**
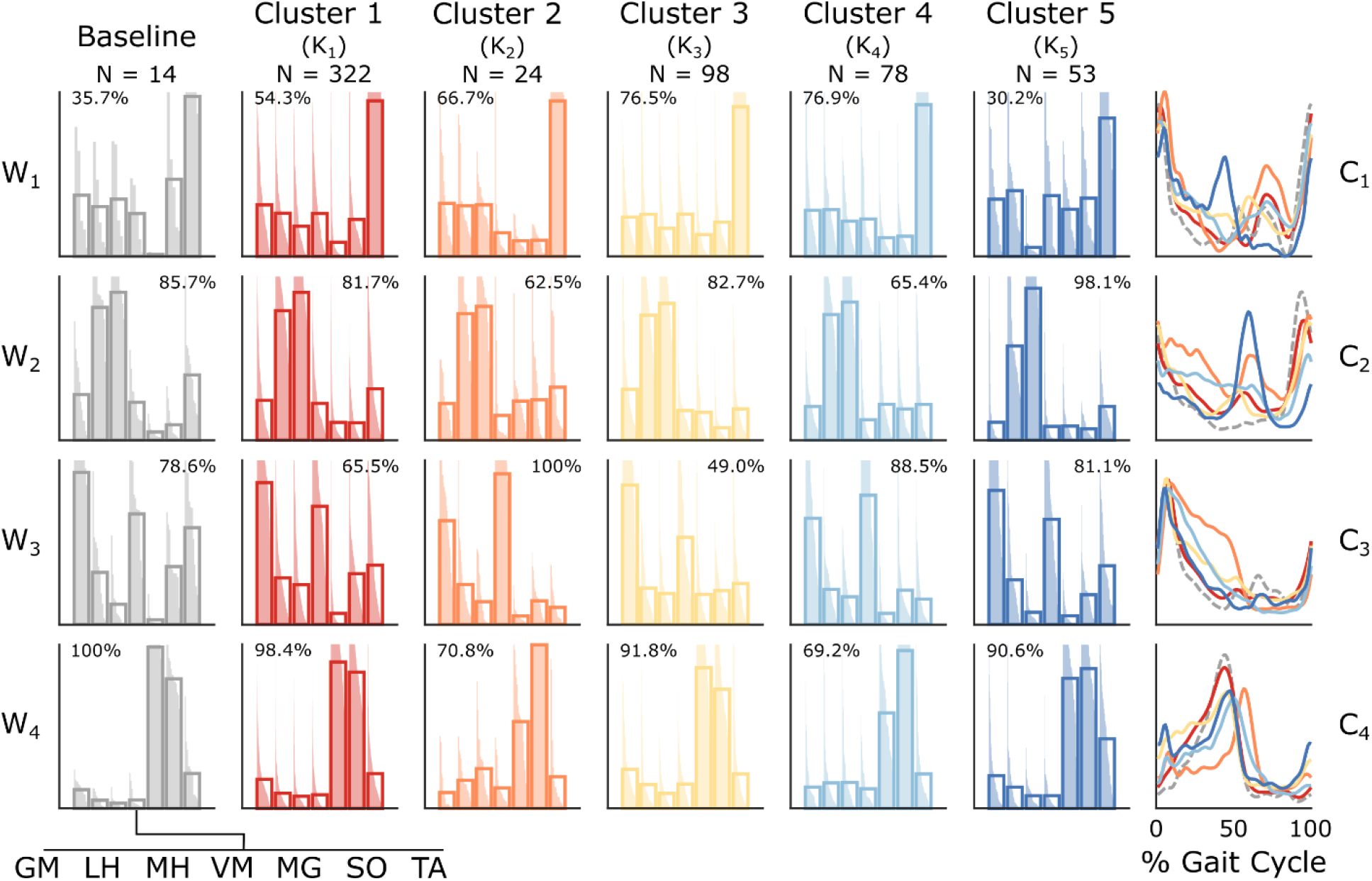
Average synergy weights (W) and activations (C) for the three-synergy solution for baseline walking and each cluster of kinematically-similar gait patterns (K_1_ to K_5_). K-means clustering was performed for the i = 3 synergy solution across all unique gait patterns and yielded four unique structures. Synergy weight plots reflect cluster-wise averages as well as weights for individual gait patterns, sorted in descending order. Percentages reflect the number of gait patterns within each cluster (i.e., K_1_ to K_5_) that used each synergy. Muscles: gluteus maximus (GM), lateral hamstring (LH), medial hamstring (MH), vastus medialis (VM), medial gastrocnemius (MG), soleus (SO), and tibialis anterior (TA).

The observed change in synergies recruited during exploration corresponded to an overall decrease in tVAF_3_BASE_ when baseline synergy weights were used to reconstruct EMG data from exploration trials (Figure 6; p << 0.001 for all groups). For the three-synergy solution, baseline synergy weights accounted for 6.0 ± 6.0% (K_1_), 17.7 ± 11.0% (K_2_), 10.6 ± 8.0% (K_3_), 15.3 ± 8.7% (K_4_), and 11.3 ± 7.1% (K_5_) less of the variance in EMG data than weights extracted directly from each unique pattern. Further, reconstruction quality was different between clusters (p << 0.001), with the largest change in synergy structure seen in K_2_. This suggests that baseline synergy weights captured muscle activity for certain gait patterns better than others, further confirming the flexible recruitment of synergies to changing biomechanical constraints.

**Figure 6:**
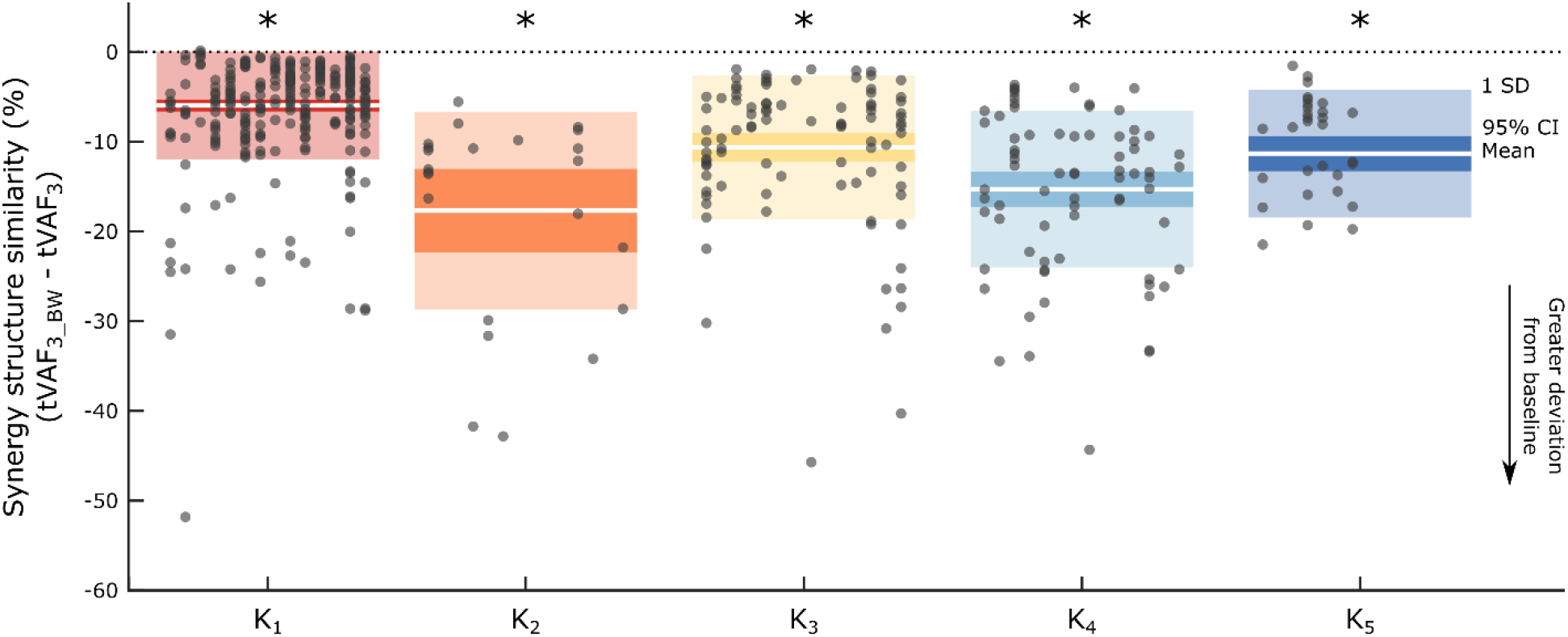
Similarity of the i = 3 synergy solution for each cluster (K_1_ to K_5_) as compared to baseline walking. Baseline synergy weights were used to reconstruct the EMG data for all unique gait patterns within each cluster. tVAF_3_BASE_ represents the amount of variance accounted for by baseline walking synergy weights and tVAF_3_ represents the variance accounted for by weights extracted directly from EMG data for each unique gait pattern. For each box, the white bars represent mean values, the solid-colored blocks represent a 95% confidence interval, and shading shows ±1 SD. Dots represent unique gait patterns and are arranged in columns to represent individual participants (P1 to P14). Large differences between tVAF_3_BASE_ and tVAF_3_ indicate that synergies during gait pattern exploration deviate more from baseline walking. * denotes significant difference between each group and zero, indicating a change in synergies from baseline walking.

### 3.3 BART Analysis

The BART model was able to explain changes in synergy complexity observed during exploration (R^2^ = 0.88; RMS error = 4.4). Baseline tVAF_1_ emerged as the top predictor of ΔtVAF_1_ (net effect = 4.6%), as individuals with higher baseline complexity increased tVAF_1_ to a greater extent during exploration than those with lower baseline complexity (Figure 8). However, this observation partially reflects the effects of regression to the mean. After baseline tVAF_1_, kinematic and kinetic predictors, especially those at the knee and ankle, had the largest effects on ΔtVAF_1_ (Figure 9A). In particular, greater knee flexion (net effect = 3.2%), anterior pelvic tilt (2.3%), hip extension moment (2.7%), and ankle dorsiflexion moment (2.8%) through stance corresponded to a greater decrease in synergy complexity. Increased knee extension moment (net effect = 3.1%) at initial contact also corresponded to less complex control. Interestingly, only one swing-phase variable had a large effect on ΔtVAF_1_; decreased knee flexion during swing resulted in greater decreases in synergy complexity (net effect = 3.3%). Further, two measures of kinematic and kinetic variability emerged among the top predictors in the BART model (Figure 9B), highlighting the sensitivity of synergy complexity to the increased stride-to-stride variability observed during gait pattern exploration.

**Figure 7:**
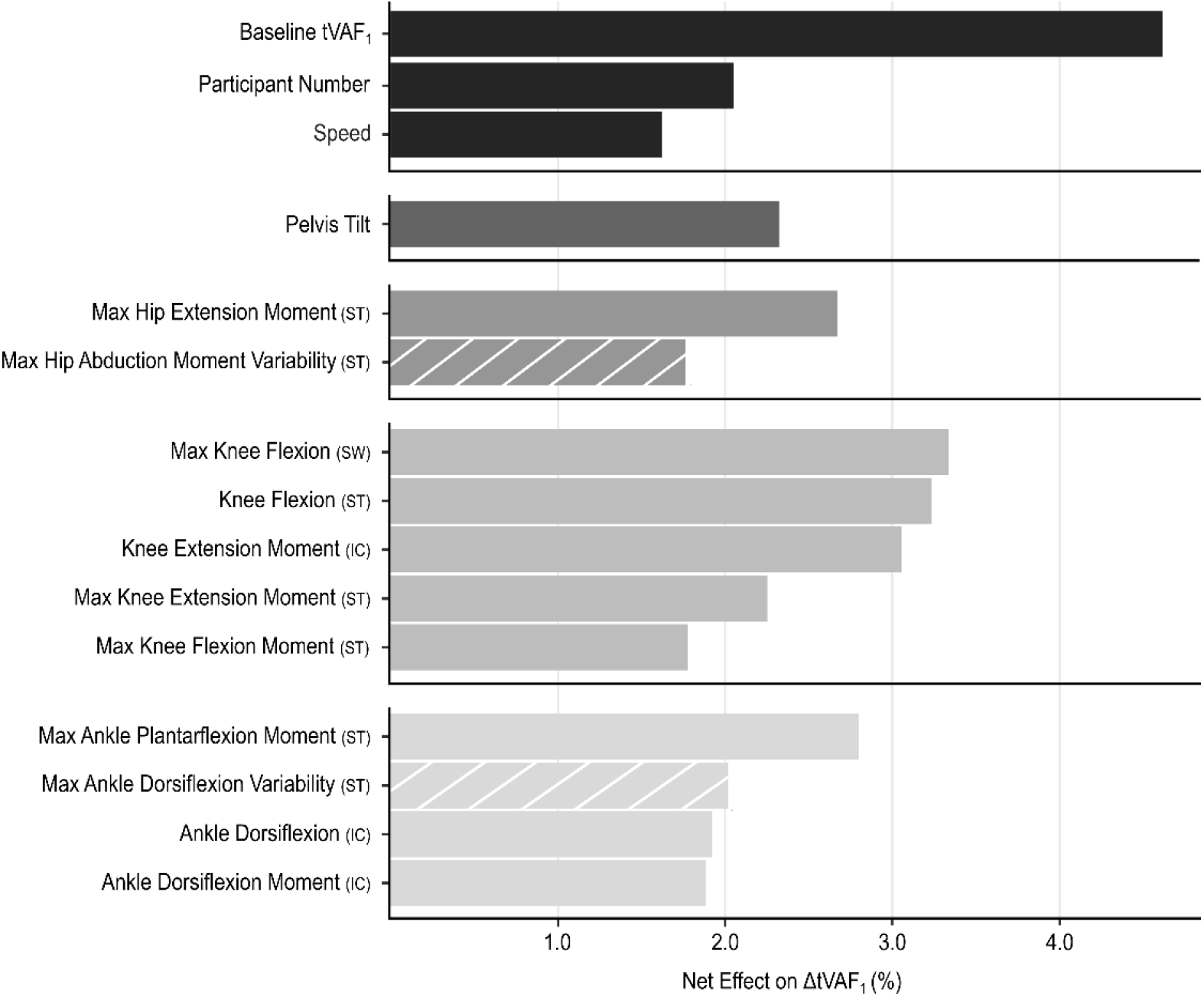
Net effects for the top fifteen predictors on ΔtVAF_1_ from the BART model. Net effects were derived from the generated ALE plots and defined as the difference between the 95^th^ and 5^th^ percentile of the response variable over the range of each predictor, when controlling for all other model covariates. Cross-hatching indicates measures of stride-to-stride variability. Gait phases: Initial contact (IC), stance (ST), and swing (SW). See Table 1 for all variables included in the BART model.

**Figure 8:**
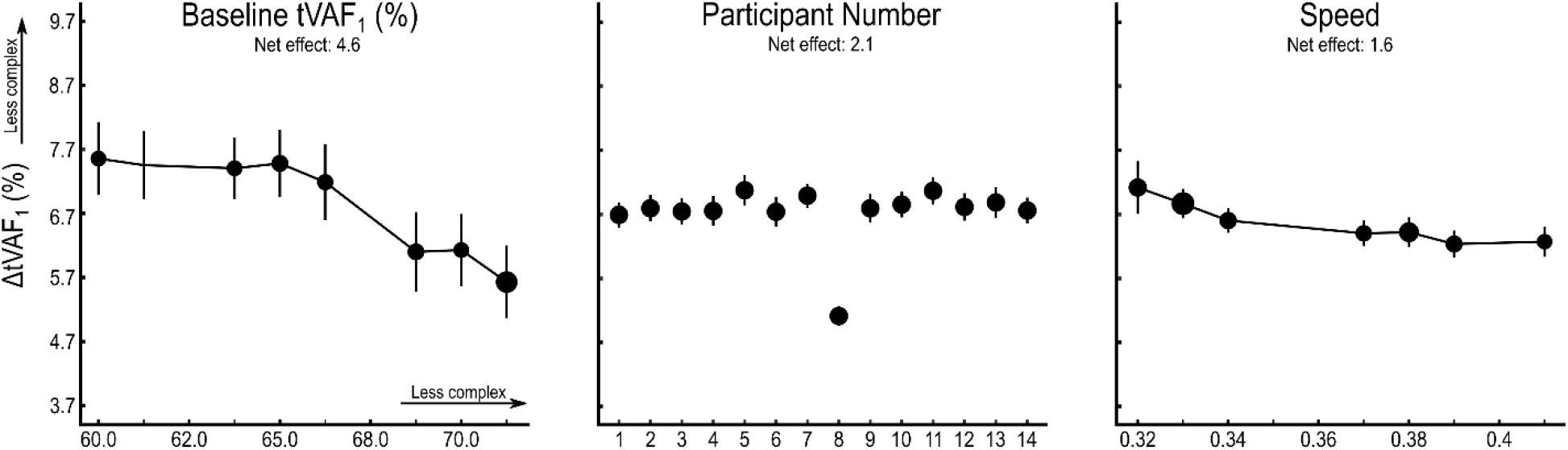
Accumulated local effect (ALE) plots for baseline synergy complexity, participant number, and speed. Speed is normalized to participant leg length. Each plot depicts the effect of an individual predictor on changes in synergy complexity from baseline, conditioned on all other predictors in the model. Predictor data is separated into evenly spaced bins and the size of individual points represents the number of samples in each bin. Larger values for tVAF_1_ indicate less complex control. Net effects were calculated as the difference between the 95^th^ and 5^th^ percentile of ΔtVAF_1_ over the range of each predictor.

**Figure 9:**
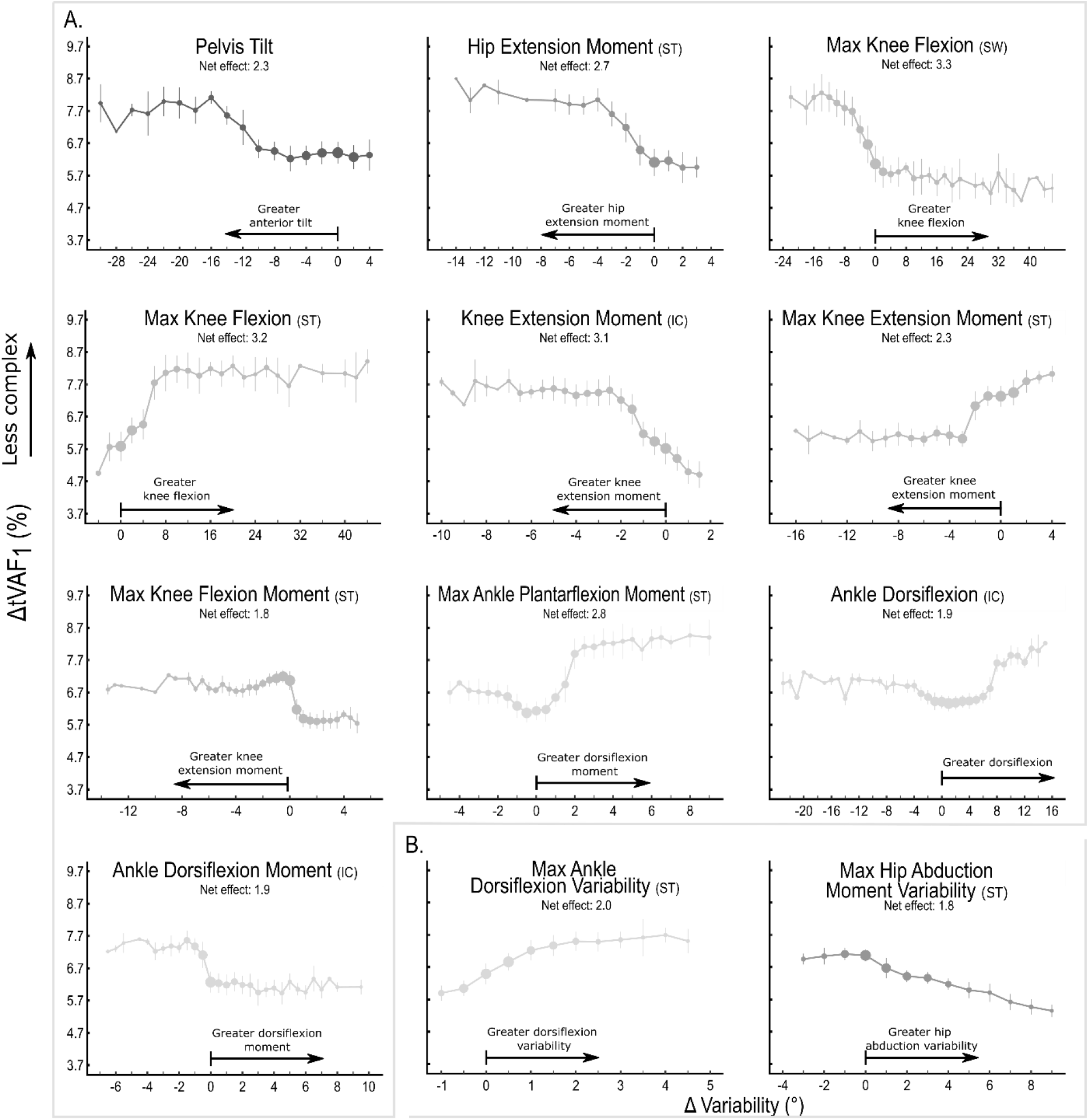
Accumulated local effects of kinematic, kinetic, and variability measures on ΔtVAF_1._ Kinematic and kinetic measures (A) are presented as z-scores normalized to baseline walking, such that the x-axis depicts standard deviations away from baseline. Variability measures (B) are presented as degrees away from baseline variability. Vertical bars indicate ± 1 SD. Plots are cropped to display the middle 95% of the predictor data to remove extreme outliers. Predictor data was separated into evenly spaced bins and the size of individual points represent the number of samples in each bin. Gait phases: Initial contact (IC), stance (ST), and swing (SW).

Beyond gait mechanics, both participant number (net effect = 2.1%) and speed (1.6%) emerged among the top predictors in the model. Although the former effect was largely driven by one participant (P8), it still indicates that differences may exist in how individuals interacted with the biofeedback system, including both the range of patterns they explored and their comprehension of the presented metric. Further, the moderate effect of speed on synergy complexity, whereby slower speeds were associated with greater decreases in complexity (Figure 8), could suggest differences in the feasibility of performing certain gait patterns at different speeds.

## 4. DISCUSSION

This study demonstrated that a small library of synergies was sufficient to characterize a broad repertoire of gait patterns attempted during biofeedback walking, and that recruitment from this library was dependent on both the type and magnitude of gait pattern deviation. Specifically, small deviations from baseline walking were generally accommodated by altering the activations of a consistent set of synergies whereas different synergies were recruited to produce larger gait changes. Participants were also able to widely modulate synergy complexity during gait pattern exploration. However, the majority of gait patterns corresponded to an increase in tVAF_1_ (i.e., decreased complexity); across all participants, only 17.4% of attempted patterns decreased tVAF_1_. Collectively, these results suggest that although synergy structures appear to be invariant, synergies can be flexibly recruited in response to changing sensory input or biomechanical constraints. This organizational strategy is advantageous for enabling rapid learning of new movement patterns and ensuring successful navigation in complex environments (Torres-Oviedo and Ting 2010, Chiel et al 2009, McKay 2007).

Our observation that a small pool of synergies emerged during gait pattern exploration aligns closely with prior literature in both animal and human models. These studies have demonstrated that synergies are consistent across a repertoire of motor outputs (Allen and Neptune, 2012; Barroso et al., 2014; Cappellini et al., 2006; Hagio et al., 2015; Ivanenko et al., 2004) and can be flexibly combined to accommodate changes in sensory input (Cheung et al., 2005; Ivanenko et al., 2004; Kargo et al., 2010; McGowan et al., 2010; Rozumalski et al., 2017) or biomechanical constraints (Krishnamoorthy et al., 2004; Nazifi et al., 2017; Torres-Oviedo and Ting, 2010). However, beyond identifying differences in synergy recruitment across movements, the nature of our protocol enabled us to understand the factors associated with these differences with greater precision. For example, we demonstrated that small deviations at the hip, knee, and ankle, as observed in K_1_, were accommodated by baseline synergies, as baseline synergy weights largely captured the variance in EMG activity during feedback walking (i.e., tVAF_3_BASE_ for K_1_). Baseline synergies were also recruited for the majority of patterns in K_5_, as the large increase in knee flexion through swing could be accommodated by altering the activation timing of the hamstring synergy (W_2_). In contrast, patterns which were defined by large deviations in sagittal plane mechanics through stance (*e*.*g*., K_2_ and K_4_), had synergy structures more dissimilar from baseline (Figure 6). A similar relationship emerged when considering synergy complexity. The results from our BART analysis demonstrated that deviations at the knee and ankle during stance largely predicted changes in tVAF_1_ during gait pattern exploration. Specifically, greater knee extension moment, ankle dorsiflexion moment, and knee flexion through stance corresponded to reduced complexity. This finding aligns with observations in clinical crouch gait in cerebral palsy, where a crouched posture places greater demand on the quadriceps to accelerate the center of mass upward and counteract gravitational force, resulting in increased coactivation of the hamstrings and quadriceps through stance and, therefore, reduced control complexity (Spomer et al., 2022; Steele et al., 2013). Hip extension moment through stance also emerged among the top predictors in the BART model, which further indicates that increased hamstring-quadricep co-contraction had a large effect on ΔtVAF_1_.

Beyond identifying those variables which were most predictive of changes in synergy recruitment, the results from the BART analysis also allowed us to capture the non-linear relationship between gait pattern deviations and synergy complexity. Specifically, a stepwise relationship consistently emerged for kinematic and kinetic predictor variables wherein tVAF_1_ was similar to baseline values up until a certain threshold, after which changes in tVAF_1_ were larger, but generally consistent. The stability of synergy complexity measures for gait patterns similar to baseline walking further confirms the propensity for the CNS to maintain a consistent control strategy to accommodate small gait deviations. Further, the plateau in ΔtVAF_1_ observed at the extremes of each gait variable suggest that bounds exist on the extent to which synergy complexity can be modulated, at least when limited to a specific muscle set.

While outcomes from the BART analysis also revealed a monotonic relationship between baseline complexity and ΔtVAF_1_, partially reflecting regression to the mean, the overwhelming majority of patterns selected during exploration increased tVAF_1_. Although these results could reflect participant comprehension of the biofeedback system and the task instructions, they may also be indicative of the underlying control strategy employed by the CNS during learning. In novel task execution, the CNS may initially assume a less complex strategy, sacrificing efficiency for stability. This hypothesis is consistent with studies demonstrating that long-term training facilitates more efficient use of neural resources (Krings et al., 2000; Picard et al., 2013) and increased supraspinal excitability (Christiansen et al., 2020; Pascual-Leone et al., 1995; Rosenkranz et al., 2007). For example, Sawers et al (2015) demonstrated that trained dancers recruited a larger number of synergies than novices during both beam and overground walking and that the synergies recruited were sparser, both of which were used to suggest that training promoted greater selective motor control. In our study, because individuals typically explored each unique gait pattern for a short bout (~10 strides) during exploration, the CNS may have had insufficient time to tune its control strategy, contributing to the observation that participants could increase, but not consistently decrease tVAF_1_ values.

Whether synergy complexity is similarly flexible and can be consistently increased following neurologic injury is largely unknown but is especially salient for informing gait rehabilitation. Individuals with central nervous system damage recruit fewer synergies than nondisabled peers (Cheung et al., 2012; Clark et al., 2010; Fox et al., 2013; Rodriguez et al., 2013; Schwartz et al., 2016; Steele et al., 2015). Further, within these populations, both synergy complexity and structure have been associated with impairment level, as those with more severe impairments have less complex control (Cheung et al., 2012; Steele et al., 2015). This is hypothesized to reflect increased reliance on spinal circuitry over supraspinal input to shape motor outputs following neurologic injury, which may reduce the overall flexibility of synergy recruitment (Leonard et al., 1991). This relationship has been demonstrated in CP, where prior literature has reported that synergies are unchanged following surgery and biofeedback training, despite both interventions yielding measurable improvements in gait (Booth et al., 2019; Shuman et al., 2019). Further, stroke survivors with less severe impairment appear to maintain the capacity to modulate synergies during locomotor training better than those with more severe impairment (Rouston et al., 2013). Understanding whether individuals with neurologic injury can consistently alter synergy complexity and improve movement, or how interventions can support sustained changes in control remain active and important areas for future investigations. While recent literature has indicated that providing richer afferent information via spinal stimulation or sensorimotor biofeedback may promote greater supraspinal involvement and, therefore, more flexible synergy recruitment during movement, studies are still ongoing (Cheng et al., 2019; Conner et al., 2021; Gad et al., 2021).

Our observation that participant number was predictive ofΔtVAF_1_ further accentuates the need to evaluate personalized responses to biofeedback. This result suggests that even when controlling for all other model covariates, including baseline complexity, interparticipant differences in response persisted. Heterogeneous response to biofeedback training has been cited previously and may stem from both individual capacity to modify the parameter targeted by biofeedback as well as system design choices (Booth et al., 2019; Huang et al., 2006; MacIntosh et al., 2019; Sigrist et al., 2013; Spencer et al., 2021; van Gelder et al., 2017). The latter likely contributed to the results observed here. Because synergy complexity is derived from multiple data streams, some participants reported feeling unsure about how specific gait changes affected the displayed metric or struggled to conceptualize what ‘more’ or ‘less’ complex gait patterns entailed, both of which likely influenced their exploration strategy. These results highlight an inherent challenge of using motor control-based biofeedback in gait training applications and presents an opportunity to explore more interpretable biofeedback metrics that can still be used to improve control patterns. For example, the output from our model suggests that providing information on joint moments to reduce hamstring-quadricep co-contraction in early stance may elicit changes in synergy complexity, although further work is needed to extend these findings to populations with neurologic injury. Our results also demonstrate the unique advantage of using non-linear function estimation techniques such as BART in order to better interpret the inherently complex and multifactorial user-system interactions present during biofeedback training to inform future system design.

### 4.1 Methodological Considerations

Although the decision to use a biofeedback system and minimal researcher coaching allowed us to capture a broader array of patterns than have been previously examined in studies of synergies in gait, there are limitations to this approach that should be considered when interpreting the results. Because we wanted participants to freely explore using the biofeedback system, we only required them to take five strides in a selected gait pattern. This meant that the novelty of the attempted patterns was likely reflected in our results, as previously described. In order to reduce this effect, we calculated synergies from the same number of strides during exploration and baseline walking (n = 5); however, it is possible that synergies may have adapted further if we had collected a larger number of strides for each unique pattern (Oliveira et al., 2014). The unstructured nature of the protocol also introduced the likelihood of observing extreme outliers, as a given gait pattern may only be attempted by a single participant. The opportunity for outliers and observed heterogeneity of participant response informed our decision to use BART as a modeling paradigm. Because BART natively constrains tree structure, it prevents data overfitting, thereby reducing the likelihood that outliers in our data set could significantly affect model outputs (Chipman et al., 2010). Finally, despite the diversity of patterns attempted, our analysis was still limited to a subset of gait patterns making it challenging to draw definitive conclusions about the relationship between biomechanical constraints and synergies. In future studies, biofeedback systems may be useful to guide users through a sample of possible walking configurations in order to develop a more comprehensive landscape of user response. Simulation paradigms, such as those employed by Kutch and Valero-Cuevas (2012), which involve systematically iterating over a range of achievable outputs, could also be a valuable compliment to the present study to provide further insight into how synergies change as a function of gait exploration. Importantly, such analyses need to be performed in both nondisabled populations and those with neurologic injury in order to understand how injury impacts one’ s ability to flexibly alter control strategies during walking.

## 5. CONCLUSION

Using motor control-based biofeedback to encourage exploration and capitalizing on non-linear machine learning methodology allowed us to identify salient features which influence how the CNS flexibly shapes control during walking. This analysis revealed that a small library of spatially invariant synergies can be flexibly recruited to produce a diverse array of motor outputs and that recruitment changes as a function of the imposed biomechanical constraints. Specifically, our results suggest that large deviations in distal joint mechanics during stance resulted in the greatest overall change in synergy recruitment from baseline walking. Further, they indicate that other participant-level factors may affect one’ s ability to modify synergy recruitment during walking, which must be considered when designing interventions to this end. Whether the recruitment flexibility observed in this study is a luxury of the unimpaired neurological system or is maintained following neurological injury is a critical next step of this work. By modeling how synergies are modulated during locomotion, we believe that this study presents both theoretical and methodological contributions towards bolstering understanding of the neural control of movement and may aid in improving interventions for individuals with neurological injury.

## Conflict of Interest

The authors declare no conflict of interest regarding the publication of this manuscript.

## Author Contributions

A.M.S., conceived and designed this research, collected and analyzed data, prepared figures, and drafted the manuscript. R.Z.Y collected and analyzed data and revised the manuscript. M.H.S. and K.M.S conceived and designed the research, interpreted results, and revised the manuscript. All authors approved the final version of the manuscript of publication.

## Acknowledgements

The authors would like to recognize Nick Baicoianu for his help developing the biofeedback system.

## Funding

This work was supported by NIH National Institute of Neurological Disorders & Stroke, R01NS091056, NIH National Center for Advancing Translational Sciences, TR002318, and NSF Graduate Research Fellowship Program, DGE-1762114.

## Notes

### Competing Interest Statement

The authors have declared no competing interest.

