## Supplemental Figure 1 for "Muscle synergies are flexibly recruited during gait pattern exploration using motor control-based biofeedback"

### S2: Kinetic Trends during Gait Pattern Exploration

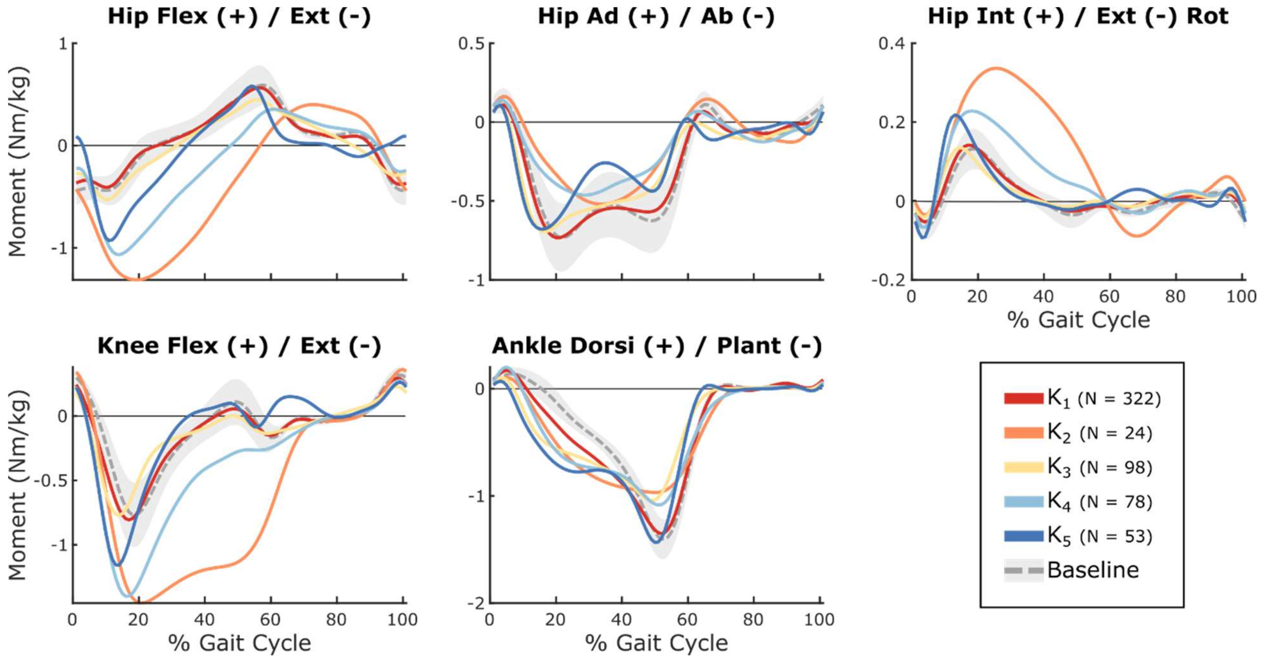


*Figure S2: Average hip, knee, and ankle kinetics for the five groups identified by k-means clustering ($K_1$ to $K_5$), representing common gait patterns attempted during exploration. The baseline condition shows ± 1SD.*
