## Supplemental Figure 2 for "Muscle synergies are flexibly recruited during gait pattern exploration using motor control-based biofeedback"

### S1: Synergy Complexity Biofeedback

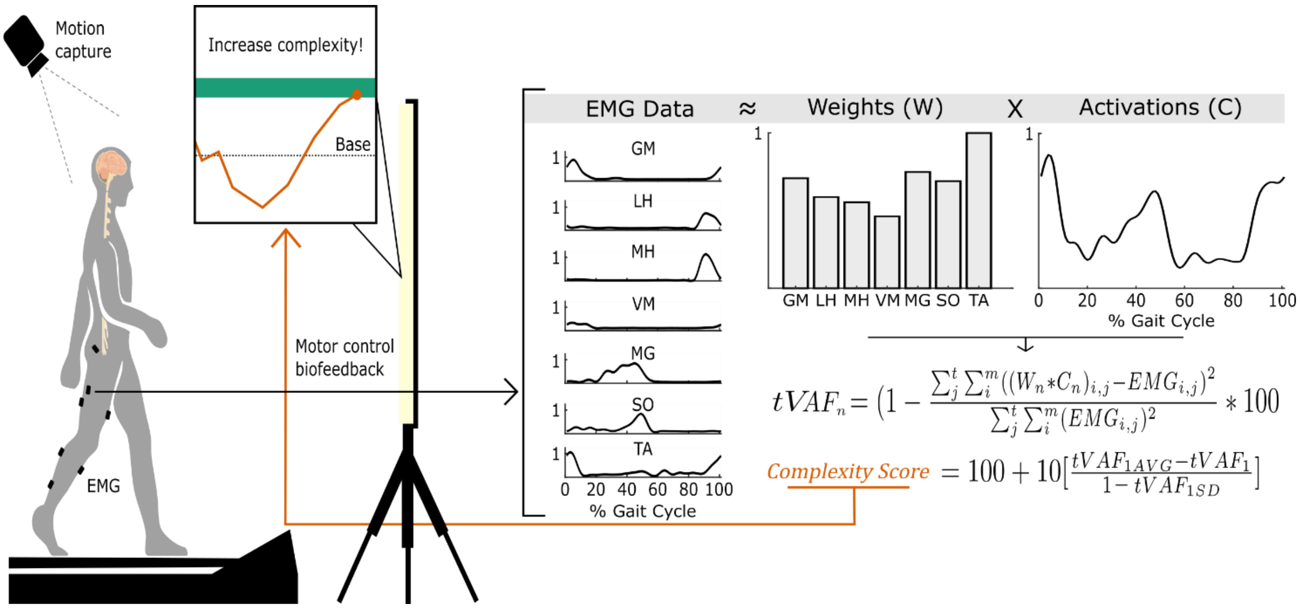


*Figure S1: Schematic of the custom biofeedback system. With each newly recorded stride, the system performed a synergy decomposition (n = 1) to calculate the one-synergy solution ($W_1$) and the corresponding activation coefficient ($C_1$), from dominant-leg EMG data for a sliding five-stride window (i.e. representing the current stride plus the previous four strides). The total variance accounted for by the one-synergy solution ($tVAF_1$) represents the reconstruction accuracy of the synergy decomposition across all muscles, m, and time points, t. $tVAF_1$ is a summary measure of motor control complexity, where a larger $tVAF_1$ indicates less complex motor control. A z-score of $tVAF_1$ is then quantified and output to the user on a visual display, alongside a green bar, indicating the goal direction of change. The complexity score is scaled such that an increase corresponds to an increase in motor control complexity. The mean ($tVAF_{1AVG}$) and standard deviation ($tVAF_{1SD}$) in the complexity score calculation are quantified for each participant from baseline data. Muscles: gluteus maximus (GM), lateral hamstrings (LH), medial hamstrings (MH), vastus medialis (VM), soleus (SO), tibialis anterior (TA), and medial gastrocnemius (MG).*
